# Adaptation at the edge: Patterns of local adaptation and genetic variation during a contemporary range expansion

**DOI:** 10.1101/2023.05.27.542577

**Authors:** Eliza I. Clark, Dan W. Bean, Ellyn V. Bitume, Amanda R. Stahlke, Paul A. Hohenlohe, Ruth A. Hufbauer

## Abstract

During range expansion, differences can evolve between populations at the core and expanding edge of a range. While theory and experimental work has focused on range expansions across uniform environments, natural range expansions often occur over environmental gradients, which present novel selection pressures. We seek to understand how genetic variation expressed in different environments may constrain adaptation during range expansion across environmental gradients, by testing whether long-established populations are better adapted to their local environments than newly established populations in the expanding edge. We study the timing of winter dormancy in a beetle introduced for biological control (*Diorhabda carinulata*), expanding from areas with cold winters to areas with milder, shorter winters. In a reciprocal environment experiment, core populations showed a pattern of local adaptation, but only some edge populations showed a similar pattern, indicating these populations vary in their degree of adaptation. Expressed genetic variation of dormancy timing in a core population was high in a local (core) environment but disappeared in a novel (edge) environment. These results show that adaptive evolution has been rapid, likely fueled by high heritability, but long-distance movement may hinder adaptation by reducing the heritable genetic variation on which selection can act.

## Introduction

Understanding the factors that allow expansions of species’ distributions is a major theme of ecology and evolutionary biology [1]. Theory and model systems show that expansion dynamics can lead to evolved differences between populations at the core of a range and those at the expanding front [2]. Most theory and experimental work on range expansion focuses on evolution of dispersal and genetic load across uniform environments, to which the populations are adapted [3,4]. However, in nature, most range expansions occur into novel or patchy environments, or across environmental gradients, such as changing climates across latitudes. A few laboratory and theoretical studies show that novel or patchy environments influence expansion dynamics [5–9], but little is known about natural range expansions over environmental gradients. Thus, a crucial area to advance research on range expansions is to understand expansion dynamics across natural environmental gradients.

Much research focuses on phenotypes expressed in populations well after range expansions have occurred, and often find evidence of adaptation to novel environments [10–13]. For example, a mosquito adapted to earlier winter onset in higher latitudes after its invasion in the United States [11,13]. In other range expansions, local adaptation to conditions such as climate and soil type has not occurred, suggesting that other factors, like phenotypic plasticity, allow persistence across a range of environments [14–16]. These studies make important advances in understanding the power of adaptation, and patterns of adaptation following successful expansion, but are generally limited in the replication of the range expansion and are conducted well after the range expansion has already successfully taken place [but see 10,11].

Thus, in addition to studying range expansions across natural environmental gradients, an important next step is to understand the evolutionary dynamics occurring during ongoing natural range expansions.

In particular, in active range expansions across environmental gradients, the process of adaptation in response to selection in novel environments will be ongoing, providing a more dynamic perspective on evolution during range expansion. If sufficient quantitative genetic variation is present, we expect that populations that have been long-established near the core of the range will be locally adapted, but that populations that are newly established may reflect their environment of origin, be in the process of adapting to their new environment, or show evidence of rapid local adaptation to the new environment.

The outcome of selection in newly established range edge populations will depend in part on the genetic variation available for adaptation, measured as heritability and evolvability. Low genetic variation will hinder the response to selection in novel environments. Since trait values, and phenotypic and additive genetic variation estimated from them, depend on the environment in which they are measured, the environment will influence the amount of genetic variation expressed in a population [17–21]. A meta-analysis on wild populations suggests that stressful environments decrease heritability, either by directly reducing the additive genetic variance of a trait, or by increasing the total phenotypic variation of a trait thereby reducing the proportion of variability explained by additive effects [22]. Across an active range expansion over an environmental gradient, we ask 1) whether populations are locally adapted to environmental conditions at the core of the range and to recently colonized environments at the edge of the range and 2) if adaptation may be constrained in novel environments by quantitative genetic variation.

We address these questions using the recent range expansion of a beetle across a latitudinal gradient from a temperate climate with long, cold winters, to a warmer environment with shorter, milder winters. A main constraint during this range expansion, and many range expansions in temperate climates, is the incidence and timing of dormancy that enables survival through the winter in cold areas but can be maladaptive in warmer areas [23]. In insects, this dormancy is called diapause. As populations move south, diapause needs to start later in the year in the newly colonized milder environment [24–26]. Our study system is particularly unique in that the introduction history of the beetle is known and the expansion has occurred along rivers, creating naturally separated expansion fronts that provide meaningful replicates. We find that populations from the core are locally adapted, while several populations at the expanding edge may be in the process of adaptation but are currently maladapted. Additionally, we find that evolutionary potential is high in core environments but is sharply reduced when core-adapted individuals are transported to edge environments, indicating that the expression of genetic variation may constrain adaptation during range expansion.

## Materials and Methods

### Study system

The northern tamarisk beetle (*Diorhabda carinulata*: Coleoptera, Chrysomelidae, hereafter tamarisk beetle), was released as a biological control agent into the United States in 2001 to help manage the invasive woody shrub tamarisk (*Tamarix spp.*). In the field, the beetle has two to six generations per year, depending on location [27,28]. Diapause in the tamarisk beetle is initiated in late summer or fall by a photoperiod cue, slightly influenced by temperature [27,29]. When reproductive individuals experience photoperiods shorter than a threshold value, they enter the process of diapause by resorbing reproductive organs and accumulating metabolic reserves within the fat body [30,31]. All life stages, from larva to adult, are sensitive to the photoperiod cue, but larvae exposed to short daylengths will continue development until the adult stage before overwintering in leaf litter beneath the host plant [30].

### Diapause timing traits

A common trait used to compare the photoperiodic cue for the timing of diapause across populations is the critical daylength for diapause induction (CDL), which is the daylength at which 50% of a population will initiate diapause [e.g., 32–34]. Daylengths longer than the critical daylength will cue diapause in fewer than 50% individuals, while daylengths shorter than the critical daylength will cue diapause in more than 50% of individuals. Critical daylength describes the photoperiod diapause cue at a population-level. However, a trait must be quantified in individuals to measure genetic variance. Here, we quantify diapause timing in individuals using the number of days a female takes to cease oviposition at a given daylength (days until diapause). Previous experiments showed that reproductive tamarisk beetles placed in diapause-inducing daylengths took between 5-20 days to cease oviposition, with fewer days required in shorter daylengths [27,30]. These results support the use of this trait as an ecologically relevant measure of the genetically encoded photoperiod cue within an individual. Days to diapause is a trait distinct from critical daylength, but importantly related, as both capture aspects of the process of photoperiod cuing of diapause [35].

### Beetle collections and lab rearing

To evaluate whether populations are locally adapted and measure quantitative genetic variation, we conducted two independent experiments with separate collections. In the test for local adaptation, we used the same collections as in Clark et al. [36]. Briefly, adult tamarisk beetles were collected by hand from *Tamarix* spp. at eight sites in the northern and southern parts of the range in the United States (**Table 1**, **Figure 1**) in Fall 2017 and Spring 2018. Northern sites were original release sites from the biocontrol program [37], and the southern sites were at the leading edge of the southward range expansion and had arrived within the previous year, except for population G (La Joya, New Mexico), which likely arrived four years prior to our collection, based on survey data (https://riversedgewest.org/documents/previous-annual-tamarisk-beetle-maps). We collected in La Joya, NM, behind the known southern distribution edge, to avoid collecting a sibling-species experiencing northward range expansion within the same river system. Beetles were reared in the lab for one generation to standardize the effects of maternal environment and females from the second lab generation were used in the local adaptation reciprocal environment experiment. For the genetic variation measurements, a separate collection was made in 2019 at site C (Delta, Utah) using the same collection methods. The second lab generation was used for the parental generation in the breeding design (see below). During lab rearing, all adults were maintained in growth chambers under reproductive conditions of light/dark 16/8hr per day and 25/20°C day/night temperatures and fed fresh tamarisk as needed.

**Figure 1.**
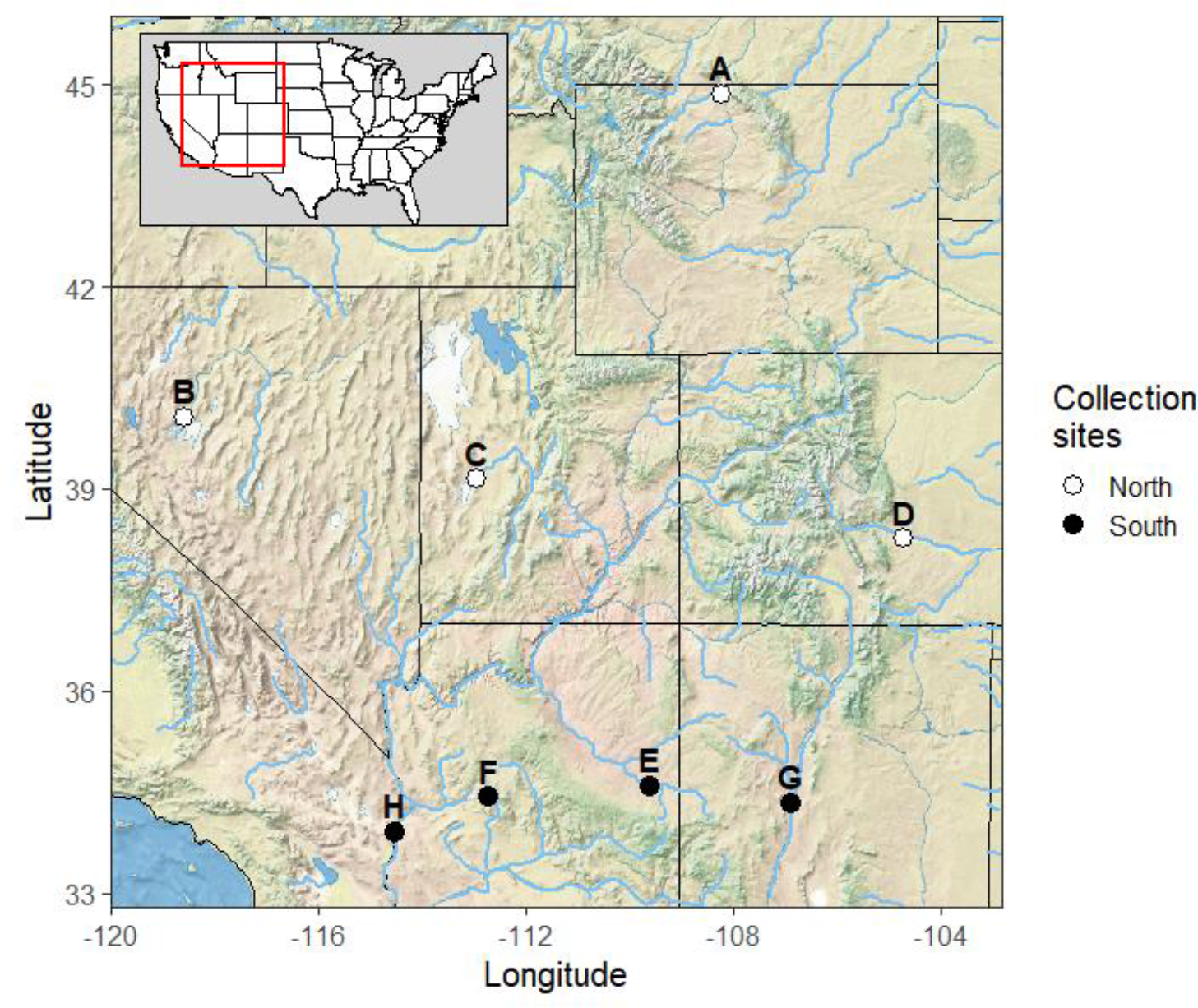
Collection sites for the tamarisk beetle in the western United States. Letters refer to **Table 1**. Northern sites are original release sites of the tamarisk beetle and are in the range core. Southern sites were at the expanding range edge when collected. Genetic variation of diapause timing was only measured for Site C. All sites were used to examine the pattern of local adaptation of diapause timing.

**Table 1.** Characteristics of tamarisk beetle collection sites. Sources and calculations of each characteristic can be found in the Supplementary Materials.

|  | Site | Lat. | Long. | Elevation (m) | Average Cumulative Degree Days | Average First Frost Date | Daylength at first frost (hrs) |
| --- | --- | --- | --- | --- | --- | --- | --- |
| North | A Lovell, Wyoming | 44.856 | -108.207 | 1115.35 | 1438 | Oct. 2 | 11.73 |
|  | B Humboldt, Nevada | 40.063 | -118.590 | 1189.62 | 1841 | Sep. 30 | 11.87 |
|  | C Delta, Utah | 39.144 | -112.958 | 1386.43 | 1767 | Oct. 8 | 11.54 |
|  | D Pueblo, Colorado | 38.268 | -104.721 | 1448.19 | 1990 | Oct. 15 | 11.27 |
| South | E Little Colorado River, Arizona | 34.593 | -109.611 | 1669.56 | 2043 | Oct. 13 | 11.45 |
|  | F Wickenburg, Arizona | 34.422 | -112.701 | 1204.74 | 3516 | Nov. 18 | 10.32 |
|  | G La Joya, New Mexico | 34.342 | -106.864 | 1433 | 2547 | Oct. 29 | 10.91 |
|  | H Blythe, California | 33.912 | -114.533 | 96.9 | 4524 | Dec. 12 | 9.94 |

### Reciprocal environment experiment to test for local adaptation

To evaluate local adaptation of individuals from the eight collection sites, we performed a reciprocal environment experiment. Mated female adult beetles were reared individually in 0.24 L plastic containers with mesh lids. Between 13 and 16 days after adult emergence, females from all populations were placed in growth chambers programmed to daylengths that represented diapause-inducing photoperiods from the north (14:20 hr:min of light per day) and south (12:40 hr:min of light per day). These daylength treatments were chosen to be near the critical daylength for diapause induction (daylength at which 50% of a population enters diapause) for Delta, Utah (site C, a northern core site), and Topock Marsh, Arizona (near site H, a southern edge site) in 2017 (unpublished data, Dan Bean). Temperatures varied between light and dark periods as in the standard rearing procedure above.

We recorded diapause incidence and days until diapause for each female in each daylength treatment above. Each day, we recorded whether each female laid any eggs in the previous 24 hours. When a female had not laid eggs in 7 consecutive days, that individual was scored as in diapause, starting from the first day with no eggs [30]. Days until diapause was measured as the number of days it took a female to stop laying eggs after switching into the daylength treatment. Females were inspected for 43 days, and females that were still laying eggs at that time were scored as reproductive (not in diapause). In our pilot experiments, there was no instance of a female that had not laid eggs for seven consecutive days starting to lay eggs again, so this was a reliable indicator that a female had started the process of entering diapause.

We can infer the fitness consequences of diapause timing in each environment based on the well-documented relationship between fitness and phenological synchrony in the tamarisk beetle [33] and many other taxa [13,24,38]. Fitness is low when diapause is either too early (which reduces the number of generations and time for reproduction, and increases predation and desiccation risk) or too late (which exposes individuals to lethal temperatures and starvation), and fitness is high at intermediate timing of and proportions in diapause. Because our treatments were chosen to be close to the critical daylength of collections from the north and south, populations that are adapted to the daylength treatment are expected to have close to 50% of individuals enter diapause. This percent is approximate because the populations are rapidly evolving [33], and limits on the number of growth chambers required that we pick only two daylengths to represent daylengths across the range. If the population is adapted to a more northern location than the daylength treatment represents (e.g., any northern population in the southern environment), more than 50% of individuals are expected to enter diapause. If the population is adapted to a more southern location than the daylength represents, less than 50% are expected to enter diapause. Given that the northern populations appear adapted with respect to diapause incidence (see results), 20-30 days to diapause appears to be an intermediate value in these experimental conditions. Days until diapause is expected to be longer than about 30 days when the photoperiod treatment is longer than the critical daylength and shorter than about 20 days when the photoperiod treatment is shorter than the critical daylength of the population.

### Measuring quantitative genetic variation

To understand how genetic variation in diapause timing and its expression may have impacted range expansion, we measured genetic variation in days until diapause and two morphological traits, body mass and thorax width. We focused on one collection site (site C, Delta, Utah) at the origin of the range expansion. Delta was an original tamarisk beetle release site in 2001 [37]. The critical daylength of populations collected from Delta has been stable for many years and it is considered to be well adapted to the location (unpublished data, Dan Bean). This site is especially relevant to the southward range expansion because it was shown to be more genetically similar to the southern sites than the other northern populations [39].

Quantitative genetic variation can be estimated by heritability (h^2^) and evolvability (I^A^) [40–42]. Heritability scales additive genetic variation by total phenotypic variation of a trait, while evolvability scales additive genetic variation by the mean value of the trait and is interpreted as the proportion change in a trait over a generation of selection. Evolvability has been suggested to be the better measure of evolutionary potential, while heritability has been used historically, so we provide both [40,42]. Because heritability and evolvability are a function of the environment in which they are measured [41], we used two daylength regimes that simulated diapause-inducing daylengths from the northern (home) and southern (away) parts of the range. The northern treatment of 13:55 hr:min of light per day was chosen as the daylength when approximately 80% of individuals from Delta would enter diapause. We chose this rather than the critical daylength, because our trait, days until diapause, requires diapause to occur to be measured. The southern treatment of 13:26 hr:min of light per day was chosen as the daylength when all or nearly all individuals from Delta would enter diapause and represents the critical daylength of tamarisk beetles living near Lake Mohave, Nevada, about 480 km south of Delta.

The temperature during these experiments was 28/20°C lights-on/lights-off.

To estimate genetic variance components of days until diapause and two additional traits of body mass and thorax width, we used a paternal half-sibling split-family breeding design [41]. Briefly, 39 sires (males) were each mated to seven or eight dams (females). Eggs were collected from each dam and reared in full-sibling families. When larvae were 3^rd^ instars, density was standardized to 15 larvae per full-sibling family per 0.24 L container, to reduce environmental variation that might obscure additive genetic variation. The three traits were measured on the adult offspring. Days until diapause was measured in the two environments described above on two mated female adult offspring per full-sibling family, one offspring per environment. Weight at adult emergence (before feeding) and thorax width were measured on the same two females and one additional male offspring per full-sibling family in a common environment that would not induce diapause (16/8 hrs light/dark, 28/20°C day/night) prior to the females entering the two daylength treatments. Realized sample size was 38 sires and 1-8 dams producing adult offspring per sire.

### Statistical Methods

All analyses were performed in R version 4.2.0 [43].

### Local adaptation

Our goal from the reciprocal environment experiment was to determine if there was a pattern of local adaptation among northern and southern collection sites in the two daylength environments. We analyzed two response variables: diapause incidence and days until diapause.

To explore the pattern of diapause incidence across the range of the tamarisk beetle, diapause incidence was predicted by beetle origin (north or south), daylength treatment (northern or southern diapause-inducing daylengths), and their interaction as fixed effects in a logistic regression model with logit link function. Collection location (factor with eight levels) was a random effect.

To examine diapause incidence at the scale of collection site and account for latitude of collection site, we fit a separate logistic model in which diapause incidence was predicted by latitude of the collection location, daylength treatment, beetle origin (north or south) and all interactions as fixed effects. 100% of individuals from four collection locations entered diapause in the southern daylength treatment, which resulted in an inability to fit these models. To resolve this, we augmented the dataset with one observation of non-diapause in each of the four impacted collection sites. We compared these results to raw means and effect sizes from Firth Penalized Logistic Regression and results were qualitatively very similar. For visualization of diapause incidence at each collection site, an additional model was fit with collection location, daylength treatment, and their interaction as fixed effects.

We ran two separate analyses on days until diapause, each including different sets of samples. First, we included all samples and all non-diapausing beetles were given a value of 43 days until diapause. Second, we excluded all non-diapausing beetles. Including all samples slightly inflates estimates of days until diapause for those that did enter diapause, but gives a more complete picture of diapause timing across a whole population, so those results are presented below. Excluding non-diapausing samples more accurately estimates days until diapause for those that did enter diapause, so results from those analyses are presented in the supplement (**Figure S1**). Conclusions about local adaptation drawn from each analysis did not differ. Responses and predictors for both sets of analyses were the same, as follows. To examine the pattern of days until diapause at a range-wide scale, days until diapause was predicted by beetle origin (north or south), daylength treatment, and their interaction as fixed effects in a Poisson model with log link function. Collection location was included as a random effect.

To examine the pattern of days until diapause at the finer scale of individual populations and understand how latitude influenced the response in each treatment, we fit a second Poisson model in which days until diapause was predicted by beetle origin (north or south), daylength treatment, latitude of collection site, and all interactions as fixed effects. Collection site means for visualization were estimated from a separate Poisson model with days until diapause predicted by collection location, daylength treatment, and their interaction as fixed effects.

All models were fit with the GLMMTMB package [44]. We visually assessed model fit with Q-Q and residual vs. fitted plots in the DHARMa package [45]. Statistical significance of effects was determined with Wald Chi-square tests in the car package [46]. Post-hoc tests of differences between marginal means and slopes of the latitude effect were done using the emmeans package [47].

### Components of genetic variation

To include non-diapausing individuals in the analysis, they were assigned a value of 43 days until diapause. We chose to include non-diapausing individuals in these analyses because they represent biologically important variation within the trait and assigning 43 days to these individuals is a conservative way to include them in the dataset. Days until diapause in the two daylength treatments were taken to be different traits and thus analyzed separately. Variance components were estimated using restricted maximum likelihood (REML). We used a half-sibling design with sire as a random effect [41]. Variance due to dam could not be estimated due to the breeding design (we assume maternal variation to be close to zero). Additive genetic variance was estimated to be VA = 4*Vsire (based on the paternal half-sibling breeding design) and total phenotypic variance VP = Vsire + Vresid [41]. Narrow-sense heritability (h^2^) was calculated as h^2^ = VA/VP. Evolvability (IA), or the expected proportional change in a trait under a unit strength of selection, was calculated as IA = VA/m^2^, where m is the trait mean [40,42].

Likelihood ratio tests were used to determine the significance of the sire variance component (Vsire). Standard error and confidence intervals around VA, VP, h^2^, and IA were calculated with a bootstrap method, following Houde & Pitcher [48]. In short, samples of sires were drawn from the observations with replacement up to the original sample size and the variance components, heritability, and evolvability were calculated as above for each sample. 1000 random samples were drawn, and variance, standard error, and confidence intervals were calculated from the resulting distribution.

For body mass at emergence and thorax width, analyses were separate for males and females, since tamarisk beetle females are generally larger than males [49]. Only one male per full-sib family was measured, so we used a half-sibling design with sire as a random effect and all variance components were calculated as above [41]. Two females per full-sib family were measured, so we used a full-sibling design with two random effects: sire and dam nested within sire. For females, additive genetic variance was estimated to be VA = 4*Vsire, as above, but total phenotypic variance was estimated as VP = Vsire + Vdam(sire) + Vresid [41]. Heritability, evolvability, and standard deviations of all variance components were calculated with the same bootstrap method.

## Results

### Local adaptation of diapause timing

For the northern populations, 57% of individuals entered diapause (95% CI 0.23, 0.85) in their local northern environment, as expected for populations well adapted to this photoperiod (**Figure 2A**). We chose a single daylength representing a northern diapause-inducing environment and expected that diapause incidence would vary by latitude of the collection location. Indeed, looking across the four populations, diapause incidence decreased with decreasing latitude at a rate of 5.23% per degree latitude (95% CI 2.06, 8.40%; **Figure 2B**). In the non-local southern environment, nearly all beetles from northern populations entered diapause (mean = 99%; 95% CI 0.91, 0.999), indicating they are maladapted to this southern light regime, as they would start diapause too early in the year (**Figure 2A**). There was no statistically clear trend among populations by latitude of origin in the southern treatment (trend = 0.20% per degree latitude, 95% CI -2.34, 2.74%; **Figure 2B**).

**Figure 2.**
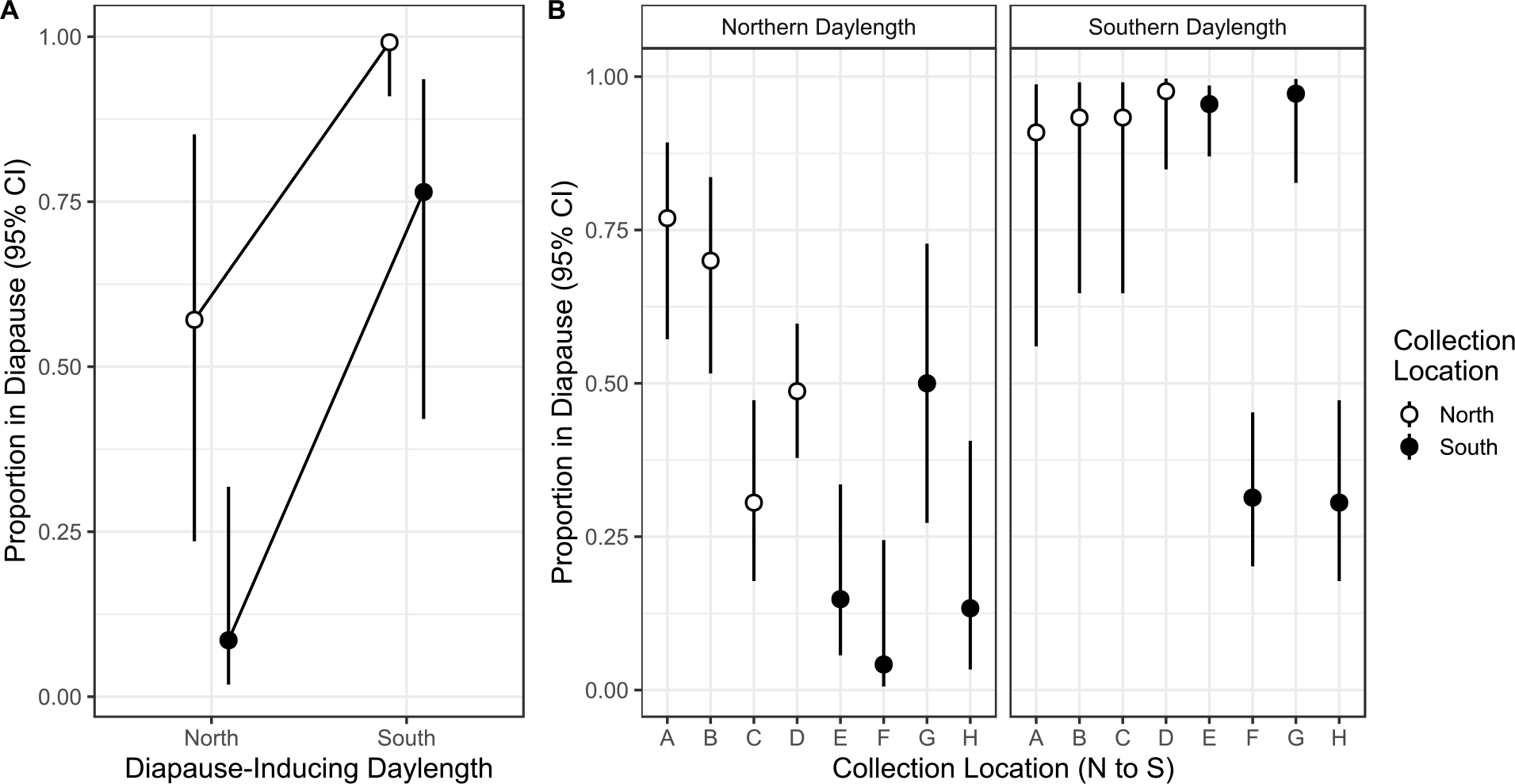
Diapause incidence in northern and southern diapause-inducing daylength treatments. Fitness of individuals will be high at about 50% diapause incidence, and low at very high or low diapause incidence. **A)** Diapause incidence averaged for northern and southern collection sites. **B)** Diapause incidence for each collection site. Collection sites are labeled as in **Table 1** and are sorted from north to south.

For the southern populations on average, 76% (95% CI 0.42, 0.94) entered diapause when in their local southern environment, indicating they are somewhat well adapted to their local light regime, since the confidence interval overlaps 50% (**Figure 2A**). When we consider variation among populations, we find that populations F and H have diapause incidences less than 50%, indicating they are adapted to a location more southern than the daylength treatment in the experiment, and populations E and G have diapause incidences above 50%, indicating they are adapted to locations more northern than our treatment represented (**Figure 2B**). In the non-local, northern daylength treatment, southern populations had on average very low diapause incidence (mean = 9%; 95% CI 0.02, 0.32), which would subject them to potentially lethal cold temperatures before they were prepared and indicating they are maladapted to this northern light regime (**Figure 2A**). Population G showed a diapause incidence close to 50% in this northern environment, indicating it may be adapted to the northern environment. For southern sites in both environments, latitude did not statistically clearly predict diapause incidence (local environment trend = 0.01% diapause/°latitude, 95% CI -0.03, 0.05%; non-local environment trend = -2.43% diapause/°latitude, 95% CI -11.95, 7.08%; **Figure 2B**).

We analyzed how quickly beetles entered diapause, or days until diapause. We included non-diapausing individuals in these analyses below to provide relative population-level estimates of diapause timing that reflect the differences in biology by latitude. Estimates for days until diapause excluding non-diapausing individuals are included in the Supplementary Materials (**Figure S1**). Northern populations in their local northern environment entered diapause in 23.8 days on average (95% CI 16.38, 34.63, **Figure 3A**). Given that northern populations appear to be well adapted to this environment with respect to diapause incidence, we take this value to be an intermediate days to diapause. Variation among the northern populations shows that the most northern populations entered diapause more quickly than other northern populations (trend = - 3.17 days/°latitude, 95% CI -3.83, -2.52, **Figure 3B**). This trend is in the direction expected, because the treatment daylength represented a date later in the year for site A than it did for site D. In the southern environment, northern beetles entered diapause in only 4.47 days on average (95% CI 3.04, 6.57, **Figure 3A**), with no statistically clear trend among populations with latitude (trend = -0.22 days/°latitude, 95% CI -0.52, 0.09, **Figure 3B**). This very short time to diapause indicates these populations are maladapted to the southern environment, since they would enter diapause too early in the year.

**Figure 3.**
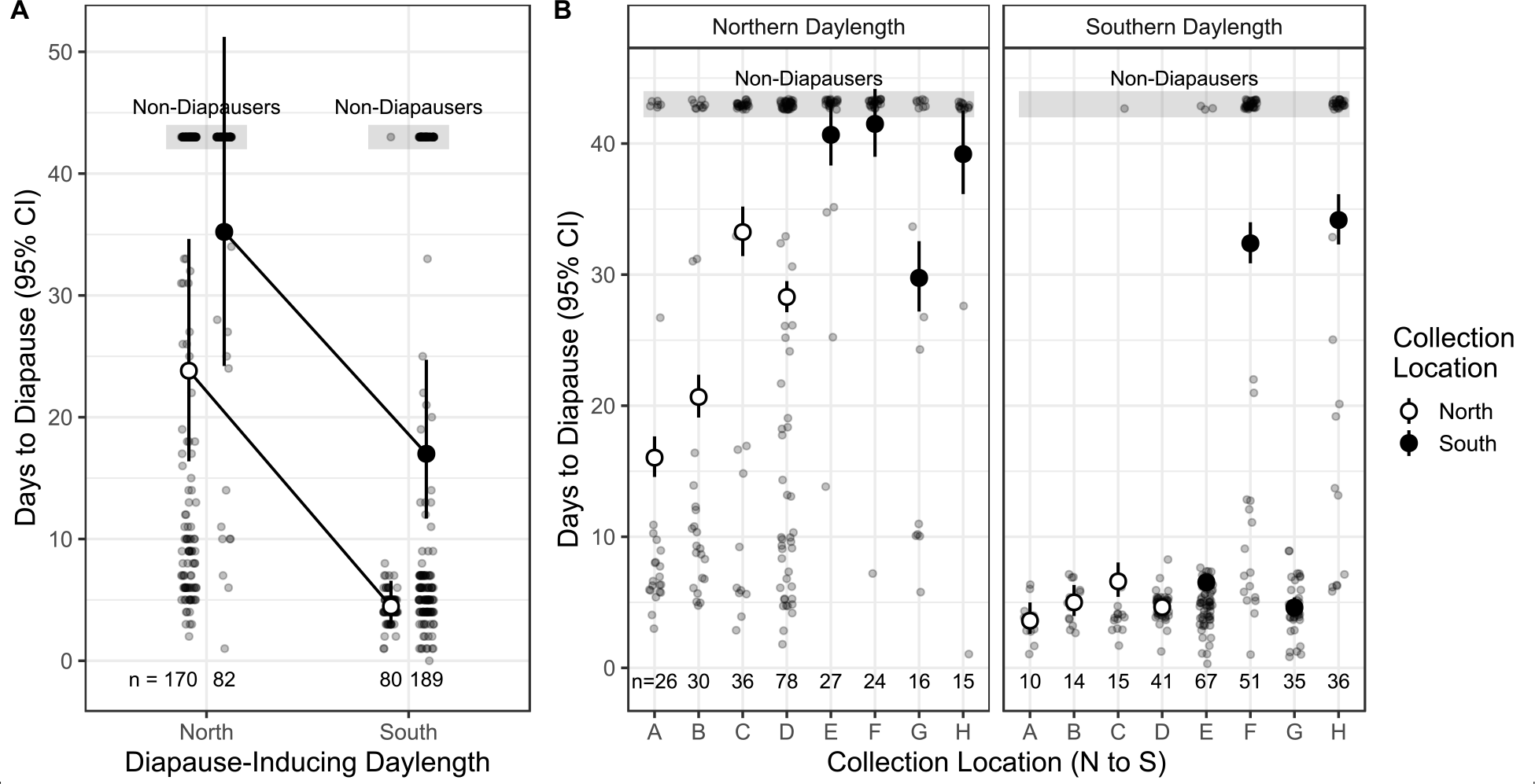
Days until diapause in northern and southern diapause-inducing daylength treatments. The numbers below points represent sample size (n). Non-diapausing individuals were included in the analyses by assigning them a diapause value of 43 days and are indicated with the grey boxes. Fitness of individuals will be highest at intermediate days until diapause, and lowest at very high or low days until diapause. **A)** Days until diapause averaged for northern and southern collection sites. **B)** Days until diapause for each collection site. Collection sites are labeled as in **Table 1** and are sorted from north to south.

Southern populations in their local southern environment entered diapause in an intermediate amount of time of 16.99 days on average (95% CI 11.68, 24.72, **Figure 3A**), which is not statistically clearly different from days to diapause for a northern population in their local environment, suggesting some adaptation to the local environment on average. However, when individual population responses are inspected, populations E and G very quickly entered diapause, indicating they are adapted to daylengths more northern than our treatment represented. Populations F and H had somewhat intermediate days until diapause, between 30 and 35 days, indicating they are adapted to the southern environment or a location slightly south of what the treatment represented. We found a statistically clear trend that days until diapause increased with decreasing latitude (trend = -0.43 days/°latitude, 95% CI -0.54, -0.32) in the local southern environment (**Figure 3B**). In the non-local northern environment, three of the four southern populations entered diapause very slowly (or not at all, **Figure 3B**), indicating maladaptation to this environment, since diapause would not happen early enough to prepare beetles for winter in the northern environment. Population G had intermediate days until diapause comparable to some northern populations, indicating that it might be adapted to the northern environment. We found no statistically clear trend with latitude among southern populations in the non-local environment (trend = 4.45 days/°latitude, 95% CI -4.55, 13.46, **Figure 3B**).

### Genetic variance of days until diapause

We estimated variance components, heritability, and evolvability of days until diapause for individuals from site C in two daylength environments. Family means for days until diapause were more variable in the northern (home) environment (10-26 days) than the southern (away) environment (7-8 days) (**Figure 4**). Genetic variance components depended on the environment in which they were measured. Variance due to sire was highly statistically significant in the northern environment (LRT1=10.93, P=0.0009), but not statistically significant in the southern environment (LRT1=0.41, P=0.52). Total phenotypic variance and additive genetic variance were reduced in the southern environment compared to the northern environment (**Figure 5A**, **Table 2**). Heritability of days until diapause was estimated at 0.76 (95% CI 0.13, 1.42) in the northern environment and 0.12 (95% CI 0, 0.52) in the southern environment (**Figure 5B**, **Table 2**). Evolvability of days until diapause was significantly positive in the northern environment (IA = 0.50), but not significantly different from zero in the southern environment (IA = 0.04, **Figure 5C**, **Table 2**).

**Figure 4.**
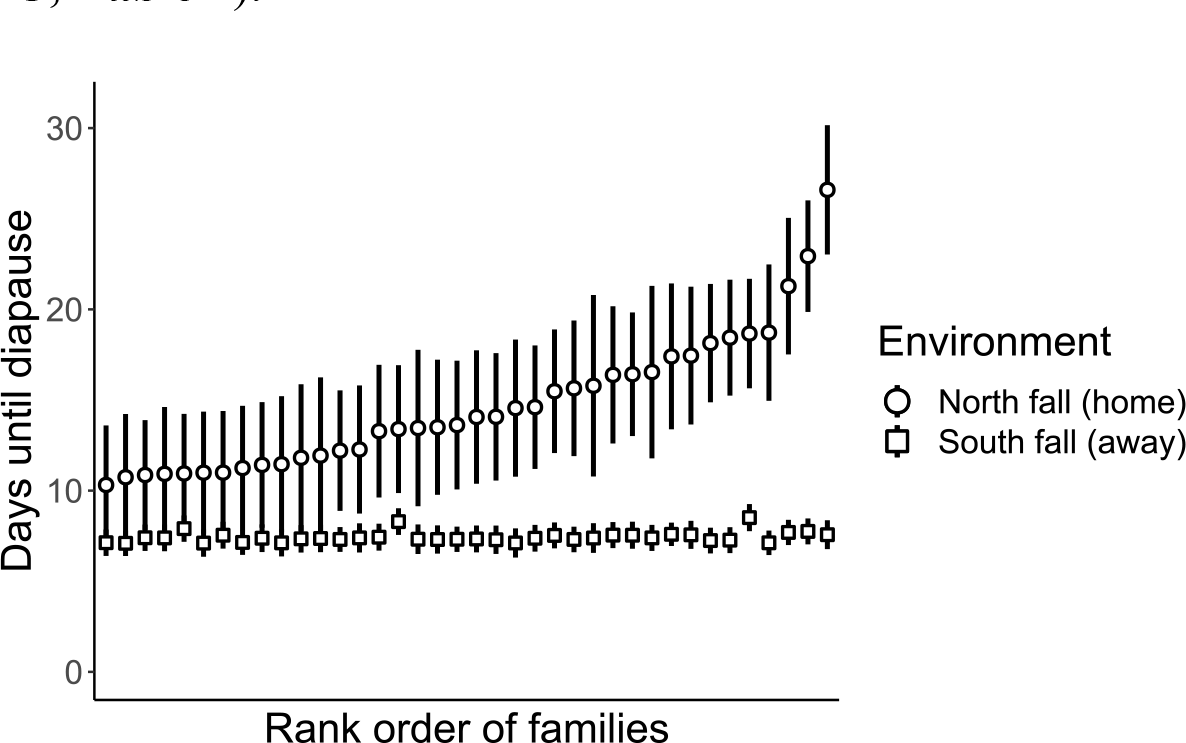
Variation in days until diapause among half-sibling families from site C in northern (home) and southern (away) daylength environments. Family estimates (+/- SD) are arranged along the x-axis in order of increasing mean days until diapause in the home treatment.

**Figure 5.**
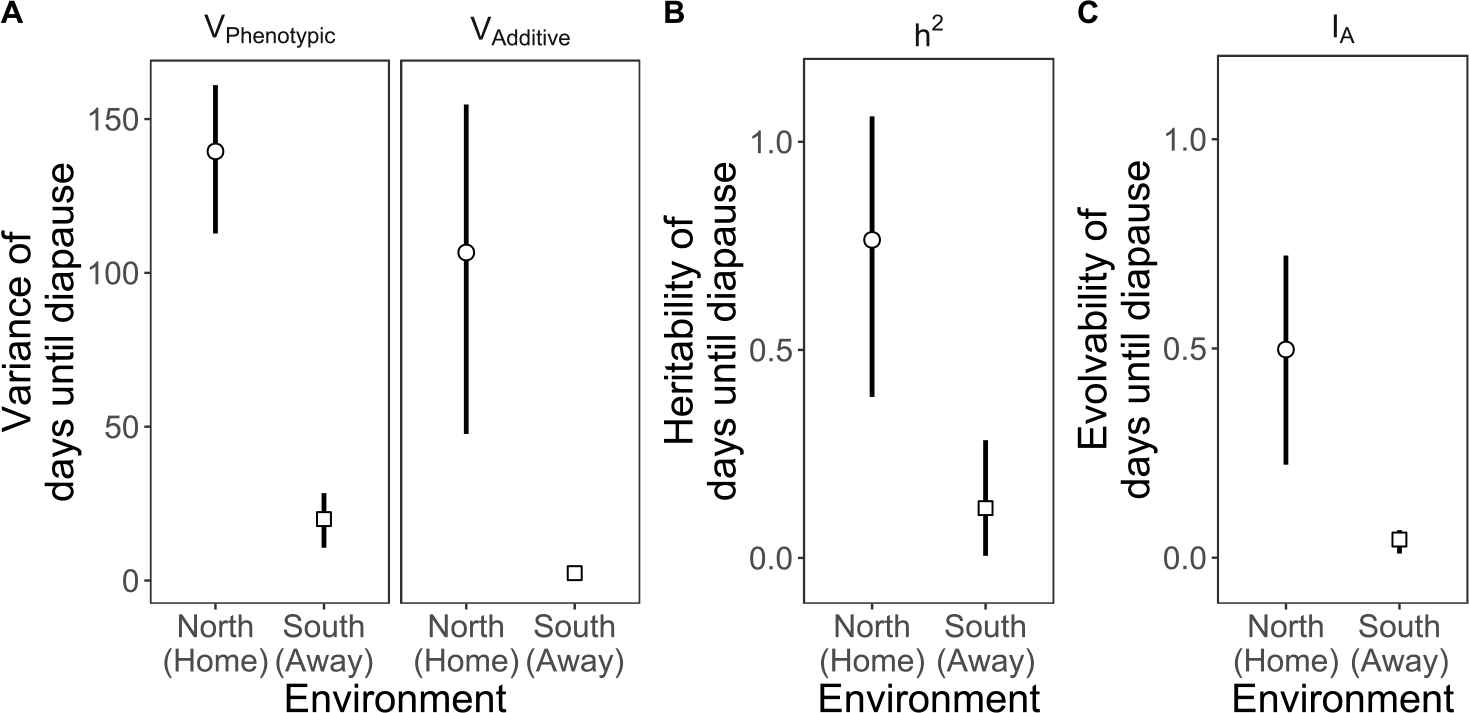
Estimates of components of genetic variation (+/- SD) for days until diapause for females from Delta, UT (site C) in two daylength environments. **A)** Total phenotypic and additive genetic variation. **B)** Heritability. **C)** Evolvability.

**Table 2.** Variance components (VAdditive and VPhenotypic), heritability (h^2^), and evolvability (IA) for days until diapause measured on females in two daylength environments. Standard deviations (SD) estimated from bootstrap procedure.

| | | $V_{\text{Additive}}$ | $V_{\text{Phenotypic}}$ | $h^2$ | $I_A$ |
| --- | --- | --- | --- | --- | --- |
| North<br>(Home) | Estimate | 107.00 | 139.00 | 0.76 | 0.50 |
|  | Bootstrap SD | 53.54 | 24.10 | 0.34 | 0.25 |
|  | Bootstrap 95% CI | (15.69, 224.03) | (92.07, 184.25) | (0.13, 1.42) | (0.07, 1.05) |
| South<br>(Away) | Estimate | 2.39 | 20.00 | 0.12 | 0.04 |
|  | Bootstrap SD | 1.54 | 8.87 | 0.14 | 0.03 |
|  | Bootstrap 95% CI | (0, 5.29) | (5.62, 39.37) | (0, 0.52) | (0, 0.10) |

### Genetic variance of body mass and thorax width

We measured the traits of body mass and thorax width on all female individuals prior to their entering the two daylength treatments, and an additional male per family. These traits provide a comparison of the heritability estimations within the same families and individuals. Body mass and thorax width are positively and statistically clearly correlated with each other (Pearson correlation=0.64). For body mass, both phenotypic variance and additive genetic variance were higher in females than males. Heritability of females was estimated to be 0.53 (95% CI 0.11-0.87) and 0.31 (95% CI 0-0.59) for males (**Figure S2**). Only the heritability estimate for females was statistically clearly greater than zero. Evolvability of body mass for both males and females were close to zero and not statistically clearly different from zero (**Table S1**). The patterns were similar for thorax width, though variance and heritability were smaller.

Both phenotypic and additive genetic variance were higher in females than males, leading to a heritability estimate of 0.36 (95% CI 0.07-0.61) for females and 0.05 (95% CI 0-0.29) for males (**Figure S3**). Only the heritability of female thorax width was statistically clearly above zero.

Evolvability of thorax width was estimated to be zero for both males and females (**Table S2**).

## Discussion

### Local adaptation

We found evidence for evolution of diapause timing in the tamarisk beetle across 10.9 degrees of latitude in the introduced range in the United States. In a striking example of rapid evolution, the newly established southern populations have diverged from the northern populations in their responses to the environments that they dispersed from only 20-30 generations (10 years) ago. Furthermore, we find that local populations respond to diapause-inducing light regimes in ways that would maintain higher fitness on average than non-local populations, suggesting local adaptation to latitude across the range expansion.

Southern sites were generally more variable in their diapause incidence and timing than the northern populations. While the four northern sites have been established since 2001 [37], the four southern sites used in this study had recently arrived at the edge of the range expansion at the time of collection. Newly colonized southern sites may require additional time to adapt more optimally to the seasonality of their locations, if genetic variation has been retained during the range expansion. Additionally, the unexpected diapause responses of the La Joya, NM population (site G) on the Rio Grande may be due to different source populations colonizing that population compared to the southern populations further east on the Colorado River and its tributaries, which could impact available genetic variation in the trait. Repeated sampling in these locations could provide additional insight to ongoing adaptation after colonization.

The variation among southern sites in this experiment may also be due to adaptation of beetles to different climate and seasonality of their local environments, such as timing and variability of winter onset and host plant senescence, and cumulative degree days. For example, the southern sites La Joya, NM and Little Colorado, AZ (E and G) appeared to be maladapted to the southern daylength regime and adapted to more northern fall daylengths, but these sites are also at higher elevations and have earlier winter onset than the other two southern sites. Indeed, in a post-hoc analysis of the southern populations in their local environment, earlier diapause (fewer days until diapause) was associated with higher latitudes and elevations, and fewer cumulative degree days. Cumulative degree days alone explained 49% of variation in diapause timing among these sites (**Figure S4**). Temperature in the fall may be one factor that is especially important in matching diapause timing with a local environment and weather in any particular year. In the tamarisk beetle, there is some evidence that diapause is delayed if temperatures are warm in the fall, especially in southern sites where winters are milder [27,29]. The role of plasticity in diapause timing in response to temperature or the quality of the host plant as food has not been well explored, but would further illuminate how adaptation and adaptive phenotypic plasticity facilitate establishment in novel environments [18,29,32]. The outcome of selection in newly established range edges will depend on how long a population has been in an area, gene flow between populations, the strength of selection and heritable genetic variation in the new environment, and the role of adaptive plasticity at each location.

We believe that the patterns we observe here are due to adaptation, rather than drift or phenotypic plasticity. An advantage of the tamarisk beetle system is its range expansion along river corridors, tracking its host plant, which provide the replicate populations used in this study. Although we were only able to simulate two environments representative of northern and southern diapause-inducing daylengths, populations in their local environments exhibit the expected trends by latitude (more northern populations enter diapause earlier and faster), which we would not expect if the pattern was generated by genetic drift. Additionally, the reciprocal rearing in common gardens experimental design eliminates the possibility that these results are due to phenotypic plasticity or environmental factors in the collection locations.

### Quantitative genetic variation

We found heritable variation in body mass and thorax width of female tamarisk beetles, which coincides well with recent evidence that female body mass increased at the edge of the tamarisk beetle range expansion, but male body mass did not [36]. We found that there is substantial heritability and evolvability of days until diapause in the Delta population (site C) at a photoperiod near its critical photoperiod, but not at a photoperiod that occurs only 13 days later in the field. This shows that there is ample underlying genetic variation in diapause related traits, but that expression of this variation depends on the environment. This provides insight into why early releases of the tamarisk beetle failed below 38°N, but were able to establish there years later. In the early releases, not only did individuals mistime diapause in the southern environment [27], but the genetic variation that could have fueled adaptation was not expressed. The high heritability we observed when individuals were close to the home environment suggests that adaptation can be rapid, if the environmental gradient is gradual or movement across it is relatively slow so that heritability and evolvability can be maintained. The tamarisk beetle’s range expansion since about 2010 was likely enabled by the maintenance of heritability of diapause timing during gradual movement southward, with few human translocations.

The trait days until diapause allowed us to track individual variation in the developmental response to photoperiod. It might also provide information needed to determine the molecular basis of dormancy in insects. Despite the importance of photoperiodism in determining the response to changing environments [23,50], the integration of photoperiod sensors, summation of photoperiod information, and hormonal signaling of diapause remain poorly understood [51].

Experimental work shows that the sum of information gained over several light cycles determines whether insects follow continuous development or switch to diapause [52,53]. This may proceed like a molecular bucket filling with a substance when days are short, with diapause initiated when the bucket fills up. Shorter days fill the bucket faster than longer days. We find that photoperiod summation varies between individuals (e.g., variation in bucket size or rate of accumulation at different photoperiods), is heritable, and evolves. This information may provide additional insight into the molecular mechanism and evolution of the photoperiodic cue that is not evident from population-level critical daylength measurements. Future genomic studies may be able to use this trait to examine the genes that underlie individual variation in diapause timing.

### Implications for range expansions

These results suggest that introductions or human translocations of biological control agents or species of conservation concern to ecologically distinct sites done with good intentions may actually hinder adaptation and establishment if genetic variation is not expressed in those novel environments. Indeed, most biological control agents fail to establish after release [54,55] and assisted migration of species facing human-caused climate change is also frequently unsuccessful [56,57]. Many hypotheses to explain these failures focus on the role of ecological interactions between the agent and biotic or abiotic features of its environment, while others emphasize the role of depleted genetic variation and lack of adaptation [54–56,58]. Our research reveals another alternative: underlying allelic variation may be adequate, but even relatively small mismatches with the environment may prevent it being expressed as heritable trait variation. Studies of heritability and evolvability of ecologically important traits in the relevant environments prior to biological control agent or assisted migration release may be beneficial in predicting both suitability for the environment and ability of an agent to adapt post-release.

Traits closely related to fitness are often expected to have lower heritability than traits less closely related to fitness, theoretically because traits related to fitness are under strong stabilizing selection [59], though this may also be caused by a related increase in phenotypic variability [42]. We found high heritability and evolvability of a diapause timing trait that should be closely related to fitness. Variable selection over time and adaptation to a variety of other environmental factors could have maintained high heritability and genetic variation in this population, even in the face of stabilizing selection on this trait [60].

The genetic paradox of invasions posits that introduced species will have low genetic variation because of small population sizes and repeated population bottlenecks [61–63]. The paradox is, then, how do invasive species persist, expand their ranges, and ultimately thrive in introduced environments? Some have argued that for many invasions, the assumptions of the paradox are faulty – quantitative genetic diversity remains high after introduction, and that variation can be selected on to match the population with its introduced environment [62,64,65]. For the tamarisk beetle in North America, additive genetic variation for an ecologically important trait, diapause timing, remains high, and can evolve quickly. Biological control agents may particularly fall into this category, especially if multiple source populations are collected and population sizes are maintained through quarantine and mass rearing phases [66].

As many species expand their ranges across environmental gradients, it is vital to understand the processes of local adaptation and maintenance of genetic variation in newly established habitat at the range edge. The evolutionary dynamics at the edge will depend upon the selective environments imposed by the landscape, including latitudinal gradients. The appropriate timing of dormancy will be crucial for many species in temperate climates to persist in novel environments. For invasive species, biological control agents, and natural populations facing rapid changes in temperature and seasonality caused by climate change, understanding patterns of local adaptation and expressed genetic variation will help to predict future movement and establishment in novel environments.

## Acknowledgements

We gratefully acknowledge undergraduate research assistants Jenna Galvin, Tori Applehans, Itai Boneh, Sam Chaney, Maria Mango, and Katie Schneider. This research was supported by a USDA Agriculture and Food Research Initiative (AFRI) grant to RAH, EVB, DWB, and PAH [grant number COLO-2016-09135], a USDA NIFA AFRI Predoctoral Fellowship to ARS [grant number 2020-67034-31888], an USDA NIFA AFRI Predoctoral Fellowship to EIC [grant number 2021-09368], and by the USDA NIFA [Hatch Project 1012868] to RAH.

## Data accessibility statement

Data and code are available at https://github.com/elizaic/Diapause-timing-2023.git and will be made available at Dryad digital repository on acceptance.

## Notes

https://github.com/elizaic/Diapause-timing-2023.git

### Competing Interest Statement

The authors have declared no competing interest.

