## Supplementary Material for "Adaptation at the edge: Patterns of local adaptation and genetic variation during a contemporary range expansion"

#### 1. Calculations of site characteristics

Elevation data are from USGS TNM Elevation Tool. Degree days were calculated by the phenology model for the tamarisk beetle at USPest.org with temperature thresholds of 11.1°C and 36.7°C, averaged for four to ten years before collections (2008-2017), depending on availability of weather station data. Average first frost day was calculated as the first day after the summer with a daily low temperature below 0°C and averaged for the same 10-year period. Daylength at first frost was calculated using standard daylength tables for each site and are presented to show how the appropriate photoperiod cue for diapause will change based on site characteristics. All temperature data were retrieved from National Oceanic and Atmospheric Administration for the closest available station to the collection location.

#### 2. Days until diapause estimates excluding non-diapausing samples

We found that individuals from both northern and southern collection sites entered diapause more rapidly in the southern environment than the northern environment. Additionally, the southern sites took longer to enter diapause than northern sites in both environments (**Figure S1A**). Specifically, in northern fall daylengths, northern populations took on average 10.64 (95% CI 8.33-13.58) days to enter diapause, which was 10.37 days faster than the southern populations, which took on average 21.01 days (95% CI 16.12-27.38). We found the same pattern in the southern fall environment, where northern populations entered diapause on average 2.77 days faster than southern populations (northern mean 4.13 days (95% CI 3.18-5.35), southern mean 6.90 days, (95% CI 5.39-8.84)) (**Figure S1A**). On average, sites in their local environment entered diapause in an intermediate amount of time (around 8-10 days), but sites in a non-local environment entered diapause either very slowly and rarely (southern populations in northern environment) or very quickly (northern populations in southern environment).

We also examined how latitude influenced days until diapause in each daylength treatment. For northern sites (A-D) in their local environment, days to diapause increased at lower latitudes (trend = -0.95 days/degree latitude (units?), 95% CI (-1.39, -0.52)), but there was no significant trend in their non-local environment (trend = -0.16 days/degree latitude, 95% CI (-0.42, 0.11)). For southern sites (E-H) in their local environment, we found a statistically clear trend that days until diapause increased at lower latitudes (trend = -0.10 days/degree latitude, 95% CI (-0.18, -0.03)), but no statistically clear trend in the non-local environment (trend = 571.43 days/degree latitude, 95% CI -891.92, 2034.78)) (**Figure S1B**). Very few individuals from southern sites entered diapause in the northern environment (between 1 and 8 per site), making these estimates less biologically meaningful and accounts for the very large standard error. In analyses including non-diapausing individuals, this trend was also statistically unclear (**Figure 3B**).

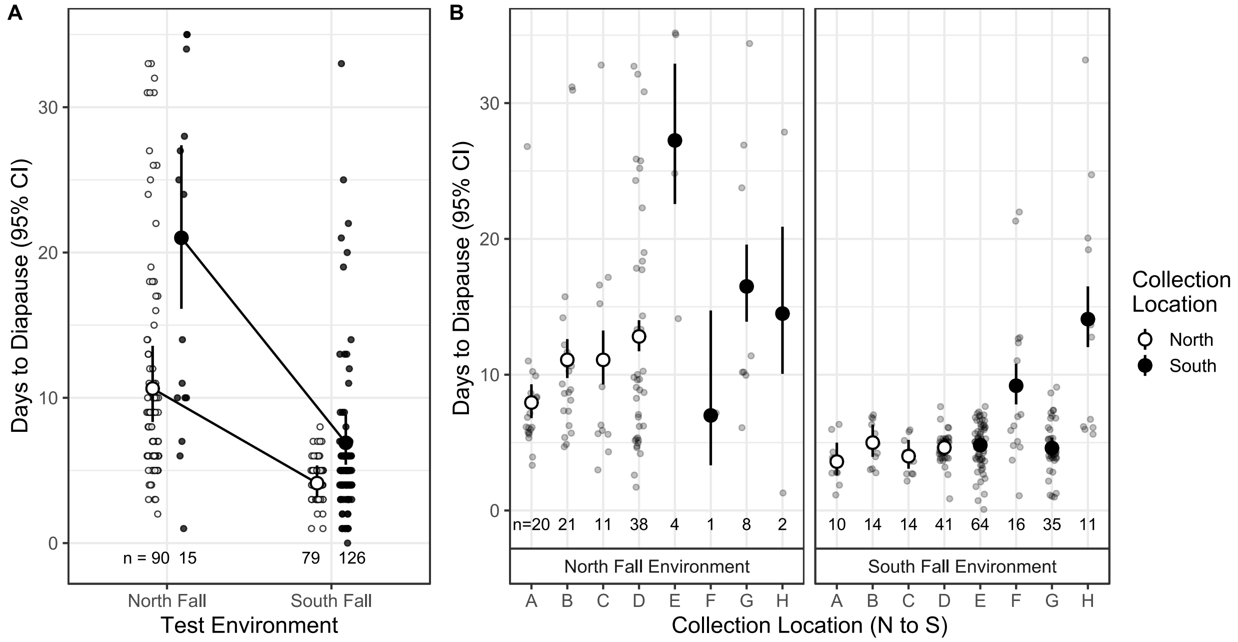


***Figure S1**. Days until diapause in daylength treatments simulating a fall environment in the north and south of the range. Non-diapausing individuals were not included in averages. The patterns shown here are qualitatively similar to patterns when the non-diapausing individuals were included (**Figure 3**), but estimates are smaller and more biologically realistic for diapausing individuals. The numbers below points represent sample size (n), which differs between groups due to unequal rates of diapause across populations. Fitness of individuals will be highest at intermediate days until diapause, and lowest at very high or low days until diapause. **A)** Days until diapause averaged for northern and southern collection sites. **B)** Days until diapause for each collection site. Collection sites are labeled as in **Table 1** and are sorted from north to south.*

#### 3. Heritability of body mass

Both phenotypic variance and additive genetic variance were reduced in males compared to females, leading to a heritability estimate of about 0.53 for females, but 0.31 for males. The sire variance component was marginally significant for males (likelihood ratio=3.553, df=1, P=0.05944), while for females it was highly significant (likelihood ratio=20.283, df=1, P=6.679e-06). Evolvability (change in trait value with unit strength of selection) of body mass for both males and females was estimated to be close to zero (stat tests).

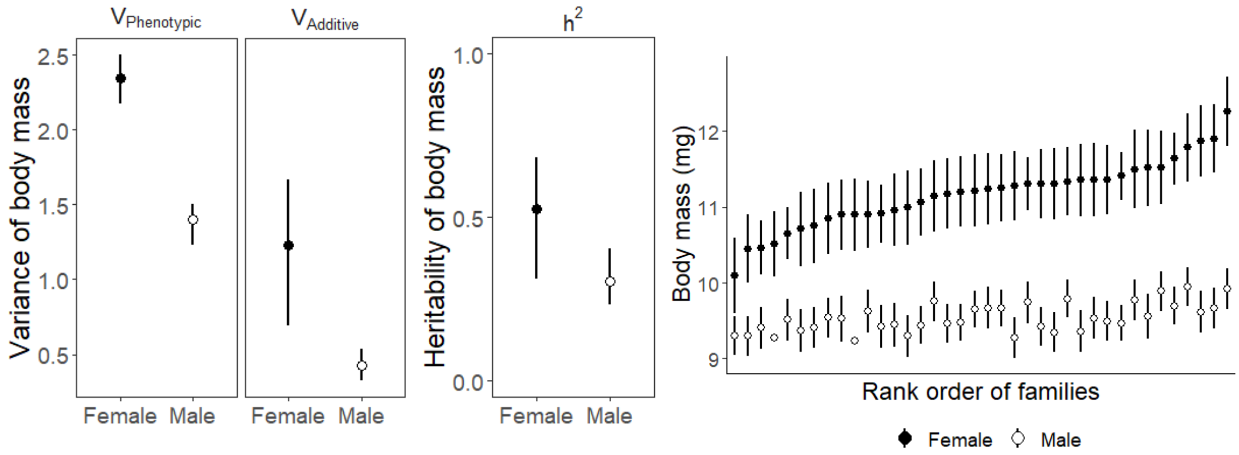


***Figure S2.*** *Total phenotypic and additive genetic variation and heritability of body mass at eclosion of males and females. The right panel shows variation in body mass for males and females by family, ordered on the x-axis by family mean of body mass for females.*

***Table S1.*** *Variance components, heritability, and evolvability for body mass of females and males. Standard deviations (SD) estimated from bootstrap procedure.*

| | | $V_{Additive}$ | $V_{Phenotypic}$ | $h^2$ | $I_A$ |
|---|---|---|---|---|---|
| **Female** | **Estimate** | 1.23 | 2.34 | 0.53 | 0.01 |
| | **Bootstrap SD** | 0.51 | 0.17 | 0.20 | 0.00 |
| | **Bootstrap 95% CI** | (0.23, 2.25) | (2.03, 2.66) | (0.11, 0.87) | (0.00, 0.02) |
| **Male** | **Estimate** | 0.43 | 1.40 | 0.31 | 0.00 |
| | **Bootstrap SD** | 0.21 | 0.12 | 0.15 | 0.00 |
| | **Bootstrap 95% CI** | (0, 0.82) | (1.16, 1.64) | (0, 0.59) | (0, 0.01) |

### 4. Heritability of thorax width

Both phenotypic and additive genetic variance were reduced in males compared to females, leading to a heritability estimate of 0.36 for females and 0.05 for males. The sire variance component was statistically significant for females (likelihood ratio=8.4198, df=1, P=0.003712), but not for males (likelihood ratio=0.13062, df=1, P=0.7178). Evolvability of thorax width for both males and females was estimated to be close to zero (stat tests).

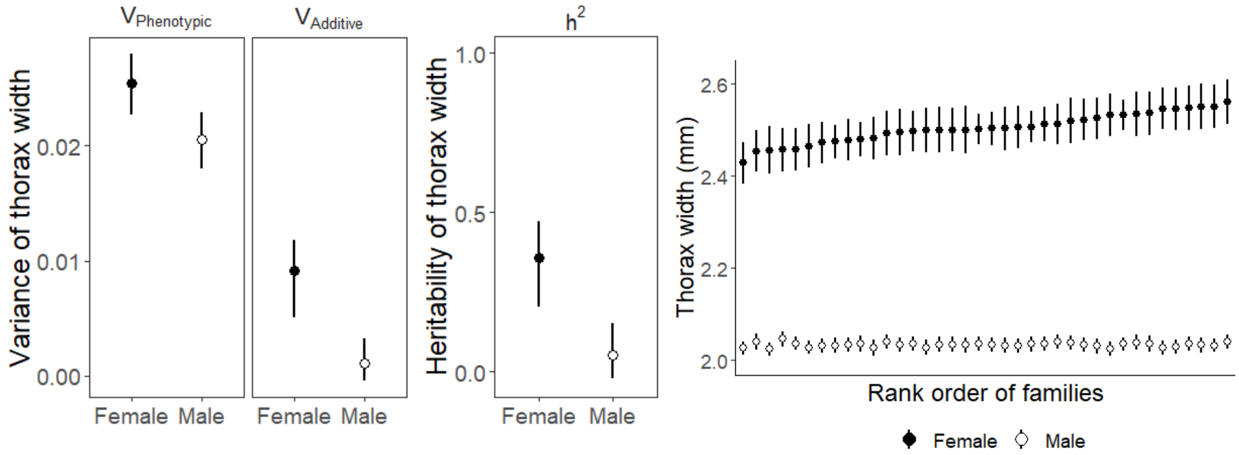


***Figure S3.*** *Total phenotypic and additive genetic variation and heritability of thorax width of males and females. The right panel shows variation in thorax width for males and females by family, ordered on the x-axis by family mean of thorax width for females.*

***Table S2****. Variance components, heritability, and evolvability for thorax width of females and males. Standard deviations (SD) estimated from bootstrap procedure.*

| | | $V_{Additive}$ | $V_{Phenotypic}$ | $h^2$ | $I_A$ |
|---|---|---|---|---|---|
| **Female** | **Estimate** | 0.01 | 0.03 | 0.36 | 0.00 |
| | **Bootstrap SD** | 0.00 | 0.00 | 0.14 | 0.00 |
| | **Bootstrap 95% CI** | (0, 0.02) | (0.02, 0.03) | (0.07, 0.61) | (0, 0) |
| **Male** | **Estimate** | 0.00 | 0.02 | 0.05 | 0.00 |
| | **Bootstrap SD** | 0.00 | 0.00 | 0.09 | 0.00 |
| | **Bootstrap 95% CI** | (0, 0.01) | (0.02, 0.03) | (0, 0.29) | (0, 0) |

#### 5. Site characteristic regressions

Diapause responses of the four southern populations used in this study were highly variable, while northern populations followed a clearer pattern based on latitude. We performed post-hoc analyses on the days until diapause response of the four southern populations in their local southern diapause-inducing daylength treatment to determine which site characteristic (latitude, elevation, or annual cumulative degree days) best predicted the diapause response.

The linear regressions predicted days until diapause with a single site characteristic for the subset of the data that was southern populations in the southern daylength treatment. In all three analyses, each predictor statistically significantly predicted the response. Earlier diapause (fewer days until diapause) was associated with higher latitudes and elevations, and fewer cumulative degree days. $R^2$ values were compared for each analysis. Latitude was the poorest of the three characteristics to predict days until diapause, while cumulative days until diapause was the best predictor for days until diapause for these populations (**Figure S4**).

These results reinforce the conclusion that populations at the southern edge of the range expansion are likely adapting diapause timing traits to several factors at the sites, including growing season length and temperature. The tamarisk beetle may be directly impacted by these factors and may also be impacted by the effects mediated by their host plant.

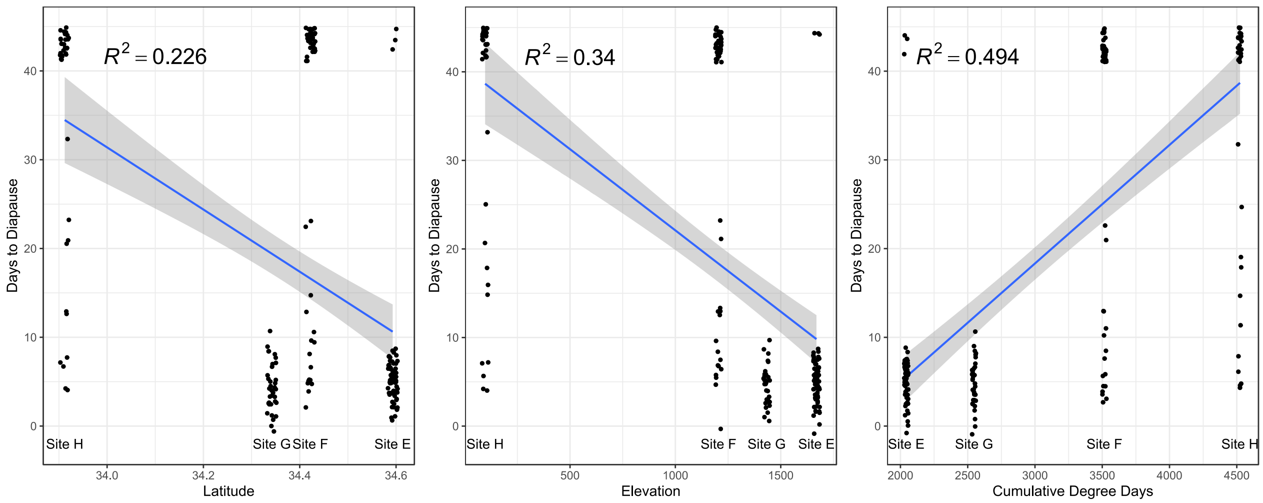


***Figure S4.*** *Linear regressions of three site characteristics—latitude, elevation, and annual cumulative degree days—with days until diapause for southern sites in the local southern diapause-inducing daylength.*
